# Neural similarity between choice options predicts group-level context effects

**DOI:** 10.1101/2025.04.15.648989

**Authors:** Asaf Madar, Tom Zemer, Ido Tavor, Dino J Levy

**Affiliations:** Sagol School of Neuroscience, Tel Aviv University, Tel Aviv, Israel 69978; Department of Anatomy and Anthropology, Faculty of Medical & Health Sciences, Tel Aviv University, Tel Aviv, Israel 69978; Coller School of Management, Tel Aviv University, Tel Aviv, Israel 69978

**Author notes:** These authors contributed equally to this work (Ido Tavor, Dino J Levy).

## Abstract

Choices are affected by the context of available alternatives, a phenomenon termed choice context effects. Current models of context effects require options to be described by two explicit numerical attributes. However, decision-makers might represent these options by additional latent attributes, which are hard to define a-priori. We propose to use participants’ neural representations to access the full attribute set they consider and predict context effects without modelling any explicit attributes. We first estimated the context effects elicited by lotteries using a behavioral sample. Then we recruited two fMRI samples with preregistered design to estimate the neural representations of each lottery without the context of choice. We predicted the context effects using only the similarity in neural representations between the individual lotteries, improving both out-of-sample and in-sample predictions compared to traditional methods. These neural representations encoded a mixture of explicit and latent attributes, previously inaccessible to researchers using only behavioral methods.

## 1. Introduction

Every decision people make in life requires them to compare between choice options that each has multiple attributes. These decisions are thought to be either based on aggregation of the options’ attributes^1^, or on applying different heuristics on these attributes^2^. Understanding multi-attribute choice behavior has been a major challenge in economics and psychology, with numerous models and theories developed over the years^3–5^. This challenge further deepens as researchers do not have access to the entire set of attributes that people represent in their minds for the choice options in question^6^.

Furthermore, it is well-known that choices are influenced not only by the attributes of each individual choice option, but also by the interaction with the attributes of other available options, a phenomenon known as choice context effects^7^. One of the most well-known context effects in decision-making is called the *decoy effect*^8^. It occurs when adding a third inferior “decoy” option to a set of two options, and while this third option is rarely selected, it influences the propensity of choosing one of the original options (“target”) over the other (“competitor”). For example, a coffee shop owner could add a medium-sized high-priced cup of coffee to a menu already including a large high-priced cup and a small low-priced cup, to increase the number of people buying the large cup instead of the small one. The decoy effect has been studied extensively over the past four decades^9–13^ using various stimuli and explained by many computational models^3,4,14,15^.

When studying decoy effects, researchers usually face two interacted problems. First, they try to understand how the attributes of choice options interact to affect participants’ choices. Second, researchers cannot describe the full attribute space of multi-attribute options, which often include latent attributes that cannot be explicitly described. For example, a car could be described only by its price and emission rating, but also has less explicit attributes such as color, overall design, and brand reputation, which affect the buyers’ choices.

Since these latent attributes are inaccessible, researchers usually describe choice options only by explicit numeric attributes in a two-dimensional attribute space, such as price and rating^9,11,13,14^. Accordingly, most theories and models require the options to have two numerical attributes in order to explain the decoy effects^8,13–15^. Hence, this two-dimensional view confines current research of the decoy effect to relatively simple stimuli, and neglects important aspects of human perception such as object representations, which are relevant for representing any object, whether in a choice scenario or not^6^. In fact, even with simple two-dimensional stimuli, participants may rely on dimensions that are hard to explicitly predefine, such as complex utility calculations and visual similarity between numbers, but nonetheless affect their choices.

In the current study, instead of relying on the researchers’ traditional two-dimensional view, we focus on the decision-makers’ view. To do so, we estimate the choice options’ representations as they are represented in the human brain using a data-driven approach, without directly modelling any pre-defined attributes. In doing so, our goal is to provide a method that generalizes to any high-dimensional choice option by integrating concepts from object representations and representational geometry into models of decision-making^16–18^.

As a proof-of-concept, we applied this approach to lottery stimuli, defined by a probability to win some amount of money (e.g., 40% of winning $20, otherwise zero). Thus, lotteries have two explicit attributes (amount and probability) and are known to elicit robust decoy effects^8,9,11–13^. We argue that even with these simple two-dimensional choice options, neural representations could capture two sources of variance of decoy effects: (1) the well-studied variance from the explicit two-dimensional attribute space (Fig 1a), (2) variance from latent attributes and positions in the high-dimensional neural space, that was previously unexamined (Fig 1d). In lotteries, these latent attributes could be the result of numerical heuristics that make some numbers easier to process^19,20^, as well as complex utility calculations and perceptual features of digits. We propose that using neural representations could provide access to both sources of variance without directly modelling any attributes, thus explaining choice context effects better.

**Figure 1.**
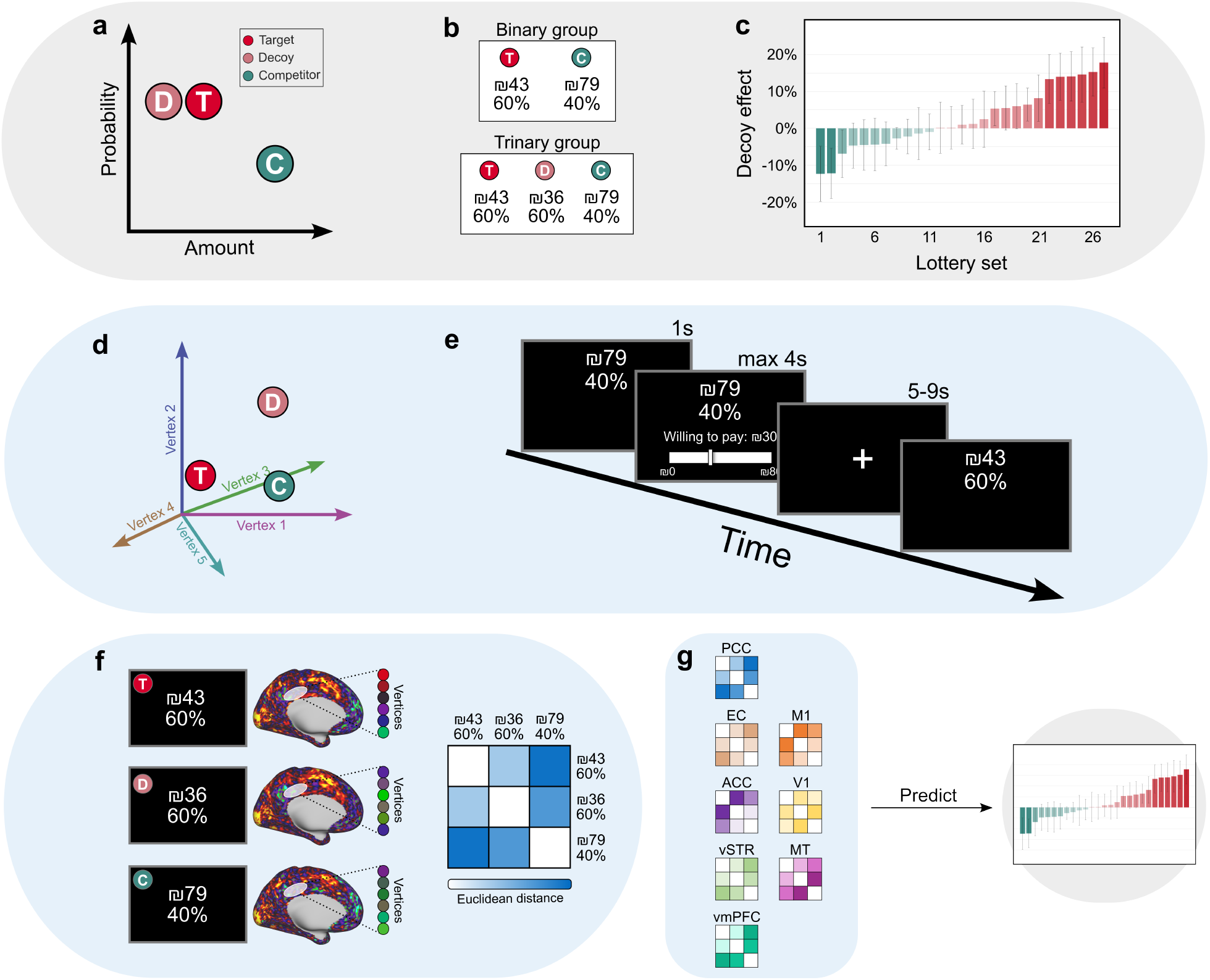
Experimental framework. (a-c) Behavioral sample experiment. (a) Traditional two-dimensional analysis of the decoy effect using the stimuli’s explicit two-dimensional attribute space of amount and probability. *T* stands for *target, D* for *decoy* and *C* for *competitor*. (b) Choice task design. Participants in the behavioral sample (*n* = 122) were randomly assigned to one of two groups: a *binary* group, choosing between two lotteries, or a *trinary* group, choosing between the same two lotteries as the binary group, but with an additional “decoy” option. In each trial, participants were asked to choose which lottery they would prefer to participate in. (c) The average magnitude of the decoy effect measured for every lottery set used in the behavioral experiment. Red bars indicate trials in which adding the decoy option increased the propensity to choose the target option and green bars indicate an increase in the propensity to choose the competitor option. Error bars represent bootstrap estimates of standard errors. (d-g) fMRI sample experiment and prediction pipeline. (d) Illustration of high-dimensional analysis of the decoy effect in the neural representation space. (e) fMRI task design. Participants in two independent fMRI samples (*n*_*first*_ = 28, *n*_*replication*_ = 34) performed a lottery evaluation task, where they viewed lotteries (1s) and then chose how much they are willing to pay in order to participate in each lottery (max 4s). The lotteries viewed in the MRI scanner were the same lotteries which constructed the lottery sets used in the choice task for the behavioral sample. (f) We first extracted a neural representation for each lottery. Then, for each participant and for each pair of lotteries of a triad, we calculated the dissimilarity between the lotteries in the neural representation space. (g) Thereafter, we averaged the representational dissimilarity matrices (RDMs) across participants for each of eight pre-defined regions of interest. Lastly, we trained models to predict the decoy effects estimated from the behavioral sample using the RDMs. PCC – posterior cingulate cortex, EC – entorhinal cortex, ACC – anterior cingulate cortex, vSTR – ventral striatum, vmPFC – ventromedial prefrontal cortex, M1 – primary motor cortex, V1 – primary visual cortex, MT – middle temporal visual area. Grey background – behavioral sample, Blue background – fMRI samples.

To test this, we first estimated the decoy context effects in a behavioral sample (*n* = 122), where participants performed a choice task between two (*binary group*) or three (*trinary group*) lottery options. The decoy effects were calculated as the difference in the propensity of choosing the *target* option between the two groups. Then, to estimate the neural representations of each unique lottery, we recruited two additional independent samples in a preregistered design (*n*_*first*_ = 28, *n*_*replication*_ = 34). These participants completed a functional magnetic resonance imaging (fMRI) scan while viewing individual lottery stimuli one by one without the context of multi-alternative choice. We then extracted participants’ high-dimensional neural representations for each lottery and calculated the similarity between all lotteries in the neural representation space. Based only on the neural representational similarity of the individual lotteries, we successfully predicted the magnitude of decoy effects as measured from the behavioral sample. We showed that both our out-of-sample predictions and data-fitting procedures surpass the performance of traditional models relying on the lotteries’ two-dimensional representations and utility estimations. Finally, we showed that the neural representations that predict the choice context effects encode information which goes beyond the two-dimensional attributes, and provided estimates to the number of dimensions encoded in each brain region.

## 2. Results

### 2.1 The addition of a third decoy option changes choice ratios

First, we aimed to measure the magnitude and direction of decoy effects at the group level for a wide range of lottery options in one sample of participants, that would later serve as prediction targets. We measured the decoy effect in a between-subjects design, which is more relevant to real-world scenarios and was widely used in previous studies^8,9,21–23^. We used lotteries since they were examined by many previous studies^9,11–13^ and they are well-suited for use in previous computational models^14,15^. Each lottery described a probability of winning an amount of money and otherwise winning nothing (e.g., 40% to win $20, 60% to win $0), with probabilities ranging from 25-75% and amounts from 4-79 NIS (≈$1 to $20). Finally, and similarly to previous studies, we constructed the lottery triads based on their relative positions in the two-dimensional attribute space of amount and probability (Fig 1a, Fig S1).

We recruited 122 participants who performed a standard decoy choice task (Fig 1b), randomly assigning them to a *binary* (*n* = 60) or *trinary* group (*n* = 62). In each trial, participants were asked to choose their preferred lottery, knowing that one trial would be eventually randomly selected and realized for a monetary prize. In the *binary group*, participants viewed two lotteries per trial, the *target* and *competitor*, while in the *trinary group* participants viewed three lotteries, the same two as in the binary group and an additional *decoy* option. They were presented with 27 unique trials. We then calculated the magnitude of the *decoy effect* for each lottery triad, defined as the difference in the propensity to choose the target option between the trinary (decoy-present) and the binary (no-decoy) groups.

Importantly, we constructed the lottery stimuli to create large variance in the decoy effect magnitudes by spanning the two-dimensional attribute space of amount and probability (Fig S1), similar to recent works^14,15^. Ultimately, our main aim was to predict this variability using neural data from an independent fMRI sample. As shown in Fig 1c, the decoy effects in the behavioral sample ranged from - 12.3% to 17.8%, with positive values indicating increased target choices due to the addition of the decoy, and negative values indicating increased competitor choices. The decoy option was rarely selected, with a median choice ratio of 2.7% (see Fig S2 for all choice probabilities). These results show that introducing a third, decoy option can shift participants’ preferences by more than 12% in either direction. Moreover, both the direction and magnitude of the decoy effect is dependent on the lottery stimuli and their combination of attributes, which change from one lottery set to another.

Finally, we tested how the options’ values affected participants’ choices by fitting a logistic regression to the binary and trinary groups separately (Supplementary Results 1). We found that in the trinary group, the probability to choose the target increased with lower target-decoy expected values (EVs) differences (*β* = −0.24, *p* < 0.001), in line with previous findings showing the target’s similarity and dominance over the decoy drives choices to the target^8^. Additionally, prior to the decoy choice task, participants performed a lottery evaluation task, in which they responded how much they are willing to pay in order to participate in each lottery. Using these willingness-to-pay (WTP) values, we found that the probability to choose the target increased with higher target-competitor WTP differences (*β*_*trinary*_ = 0.51, *β*_*binary*_ = 0.17, *ps* < 0.001), suggesting participants responded based on their previous evaluations. The difference between the lotteries’ EVs showed the same effect (*β*_*trinary*_ = 1.06, *β*_*binary*_ = 0.9, *ps* < 0.001).

### 2.2 The decoy effect is predicted by similarity of neural representations

After estimating the decoy effects from the behavioral sample, we aimed to predict them using the high-dimensional neural representation space from an independent fMRI sample. Here, instead of analyzing each lottery based on its position in the explicit two-dimensional attribute space (amount and probability), we analyzed its position in the high-dimensional neural representation space, which may include additional latent dimensions inaccessible to researchers. We hypothesized that the relative positions in this space would predict the group-level decoy effects as were measured in an independent behavioral group. Since we measured the decoy effects at the group-level using the difference in average choice ratios between the binary and trinary groups, we aimed to predict them using the average, rather than individual, neural representational similarity, measured from independent fMRI samples.

To estimate the lotteries’ positions in neural space, we ran a preregistered fMRI experiment and recruited the first sample of fMRI participants (*n* = 28). After analyzing their data, we performed a power analysis (https://aspredicted.org/s633-nkb6.pdf) and recruited the replication sample (*n* = 34). Inside the scanner, participants viewed only one lottery on each trial for a total of 31 unique lotteries and were asked to state how much they are willing to pay to participate in that lottery (Fig 1e, see Fig S3 for the fMRI behavioral results). Note that no context-dependent choices were performed during the scan. Importantly, the individual lotteries presented in the fMRI experiment were identical to the lotteries that constructed the lottery sets in the choice task of the behavioral sample. Therefore, we could use the lotteries’ positions in the high-dimensional neural space as measured by the fMRI scans to predict the magnitude of the decoy effects elicited by their combinations, as measured in the independent behavioral sample.

To do this, for each participant in the first fMRI sample we first extracted their neural representation of each lottery in eight pre-defined regions of interest (ROIs). As we aimed to take advantage of the lotteries’ representations in the latent neural space, which are not limited to the explicit attributes, we chose visual and motor ROIs, representing low-level sensorimotor information, and value-related ROIs, representing high-level information (Fig 1g; Fig S4 for anatomical locations). Then, for each participant we calculated the representational dissimilarity matrix (RDM) for each lottery triad that was presented to the independent behavioral sample (Fig 1f), including the distances between the Target and Decoy (*TD*), Target and Competitor (*TC*), and Competitor and Decoy (*CD*) lotteries. Thereafter, we averaged the RDMs across participants, resulting in three average neural dissimilarity values (*TD, TC, CD*) for each lottery triad in each ROI. We then trained an RDM-based regression based on the pre-defined ROIs’ RDMs to predict the decoy effects these lotteries elicited in the independent behavioral sample. The models were evaluated using cross-validation, such that the lottery triads in the train and validation sets were non-overlapping. Finally, we repeated the exact same procedure for the replication fMRI sample.

Using the first fMRI sample, we significantly and robustly predicted the decoy effects using only the neural similarity between the lotteries, with an average error of 6.6% between the predicted and actual decoy effects (*RMSE* = 0.0656, *p* = 0.0015; Fig 2a, b). Importantly, this model did not have direct access to the lotteries’ explicit attributes and used only the neural representations of the pre-defined ROIs including the Entorhinal cortex (EC), ventromedial prefrontal cortex (vmPFC), M1, V1, and the middle temporal visual area (MT). We replicated this result in the replication fMRI sample showing again high prediction success with an average error of 6.6% (*RMSE* = 0.0659, *p* = 0.0016; Fig 2a, b). The replication RDM model primarily used the same subset of ROIs including EC, vmPFC, M1, and MT. An analysis of the regressions’ coefficients did not show a single interpretable pattern across ROIs (Fig S5), likely because each ROI encoded a different mixture of attributes (see Fig 3-4), making it difficult to assign interpretation to individual features across ROIs.

**Figure 2.**
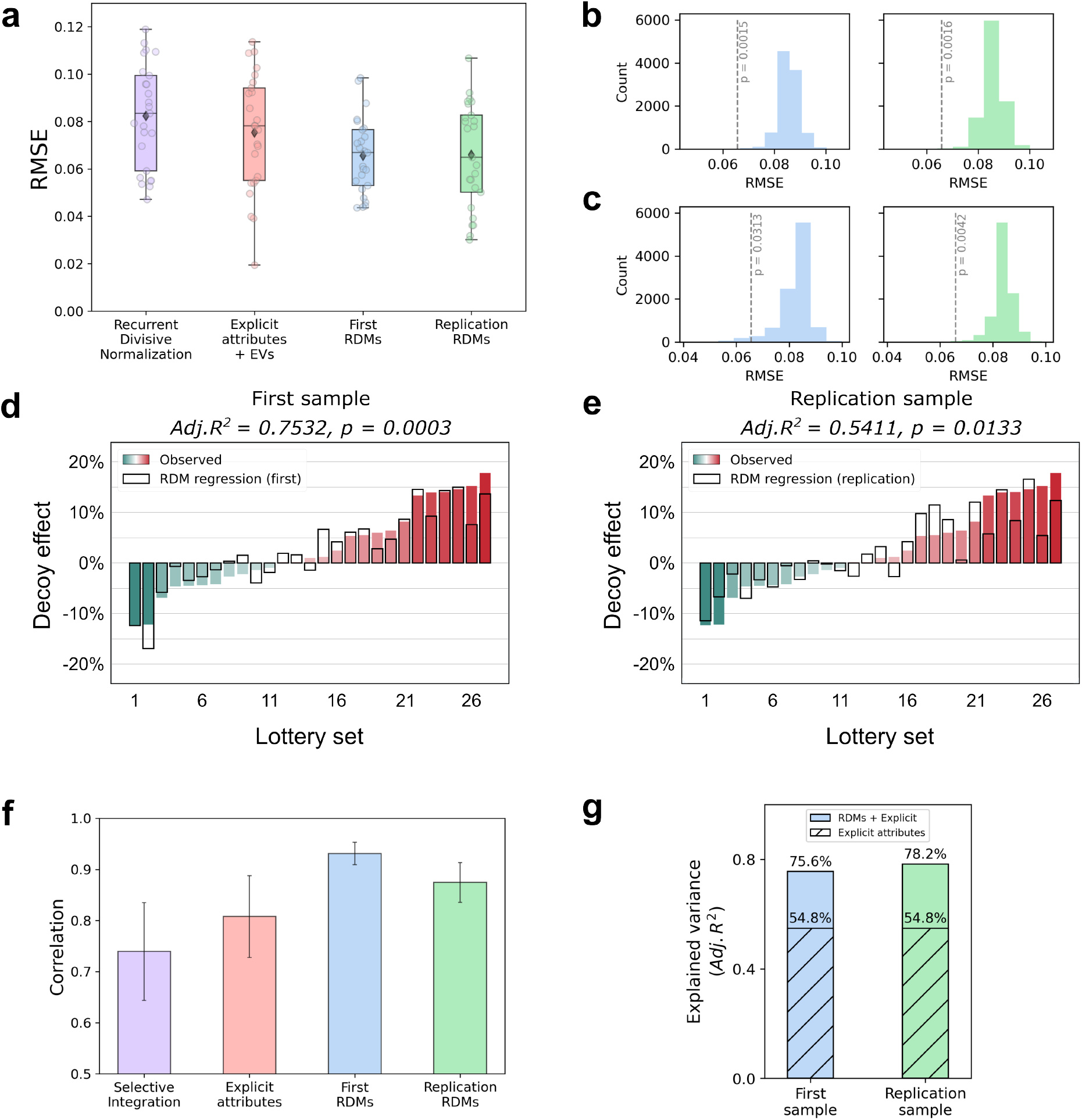
Neural representational similarity predicts the decoy effect. (a-b) Predicting the magnitudes of the decoy effects using value normalization, explicit attributes, or neural similarity. First/Replication RDM - Lasso regression models based only on the neural representational dissimilarity matrices (RDMs) between the lotteries, based on eight pre-defined ROIs. Explicit attributes model - a Lasso regression model using the lotteries’ explicit attributes (amounts and probabilities) and their expected values (EVs). Recurrent Divisive Normalization – a computational model trained to predict participants’ choices via attribute normalization^14,24^. All models were evaluated using cross-validation, such that the training and test lottery sets had no overlapping lotteries, resulting in 25 folds. Each dot represents the result of a single fold. (a) Root mean square error (RMSE) between predicted and actual decoy effects. (b) Permutation test for the RDM models’ prediction error (RMSE). Left - first RDM model, Right - replication RDM model. (c) Comparison of the RDM models’ prediction error to models that were trained on RDMs of combinations of eight randomly selected brain parcels. Left – first RDM model, Right - replication RDM model. (d-e) Data fitting procedure. Colored bars show the observed decoy effects (from Fig 1c) and black bars indicate the models’ fit. (d) The fit using the first sample, based on a forward stepwise regression model which selected the RDMs of four out of the eight pre-defined ROIs: MT, M1, EC, and PCC. (e) The fit using the replication sample, based on a linear regression model using the same four ROIs selected in the first sample. (f) Correlations between fitted and actual decoy effects for the best performing models. Selective Integration – a computational model fitted to participants’ choices using an evidence accumulation procedure with an attention mechanism^25,26^. Error bars represent bootstrap estimates of standard error. (g) Comparing the explained variance of a regression model using only the lotteries’ explicit attributes to models incorporating both the explicit attributes and the selected RDMs.

**Figure 3.**
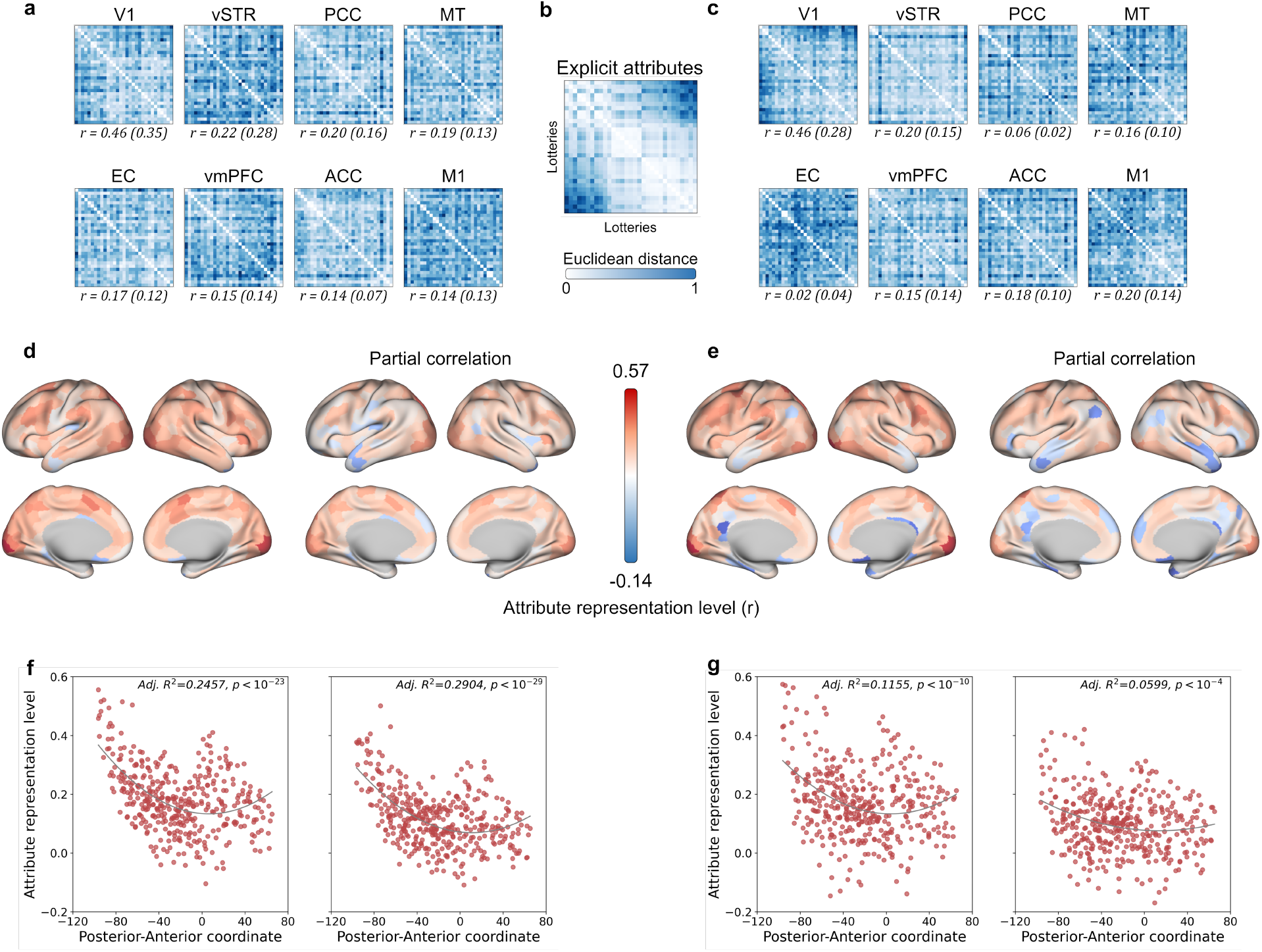
Explicit attribute representation levels. (a-c) Explicit attribute and neural space RDMs. The explicit attribute RDM was calculated based on the distances between each pair of lotteries in the two-dimensional space of amounts and probabilities. We then correlated the neural RDMs and the attribute RDM. We also calculated the partial correlations between these RDMs, after controlling for the perceptual similarity between the stimuli based on the similarity in the first layer of a convolutional neural network^30^. (a) The average neural RDMs in the first sample. ROIs are sorted by their correlation value with the explicit attribute RDM, and brackets indicate the partial correlation after controlling for perceptual similarity. (b) The explicit attribute RDM. (c) The average neural RDMs in the replication sample. The order is similar to (a). (d-e) The explicit attribute representation level across the whole brain. We correlated the neural RDM of each brain parcel with the explicit attribute RDM and projected these correlations onto a brain surface. (d) First sample, left – full correlations, right – partial correlations after controlling for perceptual similarity. (e) Replication sample, conventions the same as in (d). (f-g) The explicit attribute representation levels as a function of the anterior-posterior axis, fitted by a quadratic fit. (f) First sample, left – full correlations, right – partial correlations. (g) Replication sample, conventions the same as in (f).

We compared the RDM models to a simpler model, using the average neural response from each ROI for each lottery instead of neural similarity. In both samples, these models performed worse than the RDM models in predicting the decoy effect (*RMSE*_*first*_ = 0.0830, *RMSE*_*replication*_ = 0.0729). Importantly, however, the average neural responses in most pre-defined ROIs highly correlated with the fMRI participants’ WTP responses (Supplementary Results 4). We also repeated the RDM analysis with combinations of eight randomly selected brain parcels and found that the models based on the pre-defined ROIs performed significantly better (*p*_*first*_ = 0.0313, *p*_*replication*_ = 0.0042, *n* = 10,000 combinations; Fig 2c), demonstrating that not every eight randomly selected ROIs can be used and that the pre-defined ROIs represent unique information that helps predict the decoy effect.

Additionally, as a baseline for prediction performance, we trained two types of models to predict the decoy effect. First, we trained attribute-based regression models, using different combinations of the lotteries’ explicit attributes (amount and probability), EVs of the lotteries, and participants’ average WTP ratings. The best model, which included explicit attributes and EVs, still performed worse than the RDM models with an average error of 7.5% (*RMSE* = 0.0753, *p* = 0.0099; Fig 2a; see Supplementary Results 2 for all models’ performance). This means the RDM models improved predictions by about 13% compared to the best explicit attribute model. Second, we trained previously published computational models that fit participants’ choices based on the explicit attributes via attribute normalization^14,24^, or evidence accumulation procedures with attention^25,26^, and inhibition^26,27^ mechanisms. The best computational model was the Recurrent Divisive Normalization^14,24^ but it showed inferior predictions compared to the RDM and attributes models (*RMSE* = 0.0824; Fig 2a; see Supplementary Results 3 for all models’ performance). This shows that relying on the choice options’ positions in the high-dimensional neural space outperforms traditional models that use the explicit attributes space in out-of-sample predictions.

Next, to adhere to previous attempts investigating the decoy effect, we also used RDM regressions to fit the data and tested their in-sample predictions. Using the first fMRI sample, we trained a forward stepwise regression that significantly explained the decoy effect variance using neural similarity (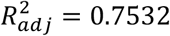, *p* = 0.0015, FDR corrected for model selection; Fig 2e), selecting four out of the eight pre-defined ROIs including MT, M1, EC, and the posterior cingulate cortex (PCC). Using the same four ROIs selected in the first sample, we replicated this result in the replication sample (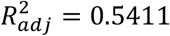, *p* = 0.0133; Fig 2f). For comparison, the best fitting attributes regression model, using only the explicit attributes, performed similarly to the replication RDM model, but worse than the first RDM model (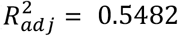, *p* = 0.0007). To compare to the best fitting computational model, which was the Selective Integration model^25,26^, we calculated the correlation between fitted and actual decoy effects for all models and found the RDM models performed best (Fig 2f; *r*_*first*_ = 0.9311, *r*_*replication*_ = 0.8677, *r*_*explicit*_ = 0.8078, *r*_*SI*_ = 0.7394). Finally, to test whether the variance explained by the RDM models extended beyond the explicit attributes, we fitted combined regression models which included the explicit attributes together with the selected RDMs (Fig 2g). Both combined models explained 20% more variance compared to the explicit attributes model performance (First: 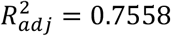, *p* = 0.0097; Replication: 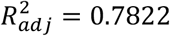, *p* = 0.0065), showing that the neural representations capture novel information beyond the explicit attributes.

Together, these results show that we can predict the decoy context effects using the positions of individual choice options in the high-dimensional neural representation space and without directly relying on the explicit attributes. We replicated these results in two independent fMRI samples, showing that similarity of high-dimensional representations predicts choice context effects in a third, independent, behavioral sample.

### 2.3 Mixture of explicit and latent attributes in neural representations predict the decoy effect

Although our models did not directly use the lotteries’ explicit attributes, the pre-defined ROIs might still represent these attributes indirectly to a large extent. To explore this, we estimated the level of explicit attribute representation in each ROI. We first calculated the explicit attributes RDM of all lotteries in the two-dimensional attribute space based on the Euclidean distance between the amounts and probabilities of each lottery pair (Fig 3b). Then, we correlated the lower triangles of this attribute-RDM and the neural RDM calculated for each ROI. The correlation magnitude indicates the extent to which each ROI represents the lotteries’ explicit attributes (attribute representation level; Fig 3a-c). Additionally, we qualitatively compared between the lottery triad configurations in the explicit attribute space (Fig S1) and in the neural space, based on multidimensional scaling (MDS).

As apparent from Fig 3a-c, all pre-defined ROIs, except V1, had low explicit attribute representation levels (*r* = 0.0162 − 0.2211), while V1 had high attribute representation levels (*r*_*first*_ = 0.4560, *r*_*replication*_ = 0.4614) suggesting that V1 largely represented the explicit attributes while the other ROIs mainly did not. Notably, the high explicit attribute representations levels in V1 were not driven solely by low-level perceptual representations, as confirmed by repeating the analysis with the partial correlations between the attribute and neural RDMs after controlling for the perceptual similarity between the lotteries (*r*_*first*_ = 0.3451, *r*_*replication*_ = 0.2752, highest level of all areas; Fig 3a-c). Finally, MDS visualizations did not reveal consistent alignment between the neural and explicit attribute space configurations in any ROI, suggesting that the representational geometry used for prediction was more complex than the two-dimensional attribute space (Fig S6).

Importantly, despite most pre-defined ROIs showing low attribute representation levels, the RDM models used their representations to significantly predict the decoy effects. This suggests that they hold relevant information for predicting choice behavior. Furthermore, most pre-defined ROIs also encoded subjective value signals from the WTP task participants performed, as evident by high correlations between the average neural responses and WTP ratings (e.g., *r*_*ACC*_ = 0.5, *r*_*vSTR*_ = 0.77, see Supplementary Results 4). Together, these findings suggest that while the ROIs did not mainly encode for the lotteries’ explicit attributes, they integrated other task-relevant representations, possibly from latent attribute spaces.

Following the discrepancy between explicit attribute representations and prediction performance in the pre-defined ROIs, we explored this phenomenon in the whole brain. Here, instead of using only the eight pre-defined sensorimotor and value-related ROIs, we used a whole-brain parcellation of 419 areas^28,29^ and calculated their RDMs exactly as we did for the pre-defined ROIs. Then, we correlated each area’s RDM with the attribute-RDM to estimate its explicit attribute representation level (Fig 3d-e). Remarkably, the resulting whole-brain map of attribute representation levels showed a posterior-anterior U-shape gradient: the highest levels are observed in posterior occipital regions, which decrease along the anterior axis, and slightly increase in anterior prefrontal regions (Fig 3f-g).

Next, we tested whether brain parcels with high explicit attribute representation levels also predicts the decoy effects better. We randomly chose eight brain parcels and, similarly to the pre-defined ROIs, trained a neural RDM-regression for each combination, and evaluated its performance in predicting the decoy effects. We repeated this procedure 10,000 times. Then, we also calculated the average attribute representation levels of each eight brain parcels. Lastly, we calculated the robust correlation between the average attribute representation levels of the eight brain parcels with the models’ prediction errors (RMSE) across the combinations. We found a weak positive relationship between the attribute representation level and prediction error in both samples (*r*_*first*_ = 0.0866, *r*_*replication*_ = 0.0745, *p* < 10^−13^), indicating that higher average attribute representation levels led to slightly worse predictions. Importantly, the small effect size suggests that predicting the decoy effect from neural representations does not rely on either explicit or latent attributes alone. Together with the pre-defined ROIs’ results, this strengthens the notion that a mixture of both latent and explicit attributes encoded in the neural representations supports our predictions.

### 2.4 Low explicit attribute representation is tied to higher effective dimensionality

After showing the decoy effect is predicted by areas that do not necessarily represent the two-dimensional attribute space, we sought to explore these other, possibly latent, dimensions. For example, area MT, which contributed strongly to the predictions, has a relatively low explicit attribute representation level (*r*_*first*_ = 0.1868). Thus, this could reflect a high-dimensional space, where only a few dimensions correspond to the explicit attributes, leading to an overall low explicit attribute representation level. However, it is also possible that MT represents a low-dimensional space of latent attributes which have low correspondence with the explicit attributes. More generally, each ROI can show four possible patterns in its representations: low or high explicit attribute representation levels represented in low- or high-dimensional spaces. In this section, we aimed to disentangle these possibilities.

To this end, we used the effective dimensionality metric, which is a continuous measure quantifying the unique information represented in a set of measurements^31,32^ (Fig 4a-c). High values indicate high dimensionality, where information is distributed more uniformly to all measurements (vertices), while low values indicate low dimensionality, where most unique information is concentrated in a small set of measurements. Together with the results of the explicit attribute representation levels from the previous section, we could characterize each ROI by these two metrics.

**Figure 4.**
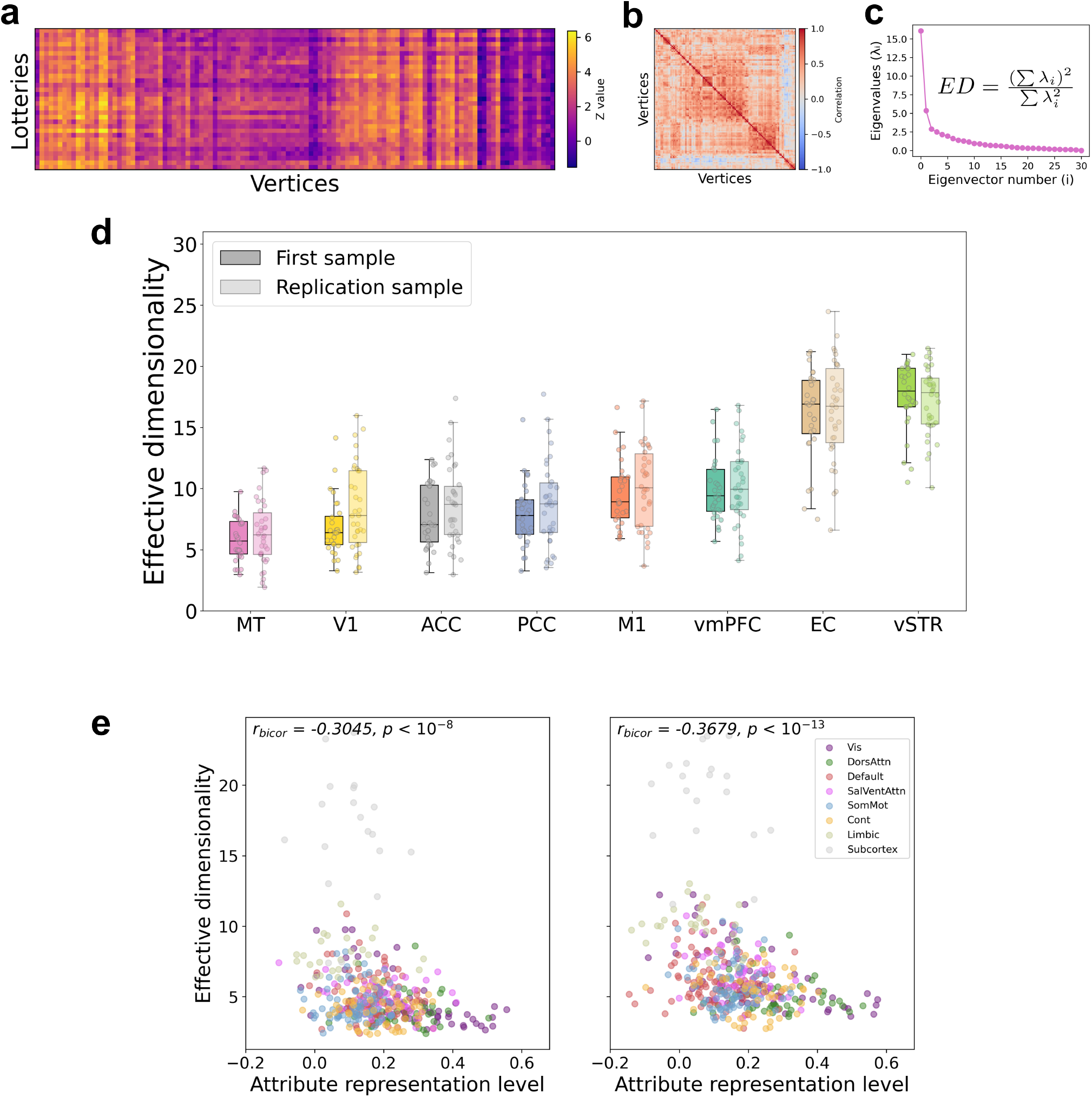
Effective dimensionality analysis. (a-c) Calculating effective dimensionality. (a) Example of a participant’s neural representations of all 31 lotteries from area MT (114 vertices). (b) The correlation matrix showing the dependency between the vertices of area MT. (c) The effective dimensionality metric is a continuous measure of the amount of unique information within a set of vertices, calculated based on the eigenvalues of the vertices’ covariance matrix. (d) The effective dimensionality of each ROI’s representations. ROIs are sorted by the average effective dimensionality across participants. The order is highly preserved across the two samples (*r* = 0.99). EC and vSTR are significantly larger than all other ROIs in both samples (*ps* < 10^-8^, FDR corrected). For the full set of pairwise comparisons see Fig S3. Each dot represents the effective dimensionality of one participant. The line represents the median. (e) Whole brain analysis. The robust correlation (biweight midcorrelation) between each brain parcel’s explicit attribute representation level and its average effective dimensionality. Parcels are colored by the canonical seven functional networks and subcortical regions^28,29^. Left – first sample, Right – replication sample.

We calculated the effective dimensionality for each participant and for each ROI, and found that visual areas (V1 and MT) had the lowest average effective dimensionality, while value-related areas (EC and vSTR) had the highest (Fig 4d). This order of dimensionality was highly preserved across both fMRI samples (*r* = 0.99). Pairwise comparisons revealed that EC and vSTR had significantly higher effective dimensionality than all other areas (*ps* < 10^−8^ in both samples, FDR corrected), while V1, ACC, and PCC did not differ from each other (*ps* > 0.1298; For all pairwise comparisons see Fig S7).

Interestingly, despite having similar effective dimensionality, V1, ACC, and PCC had very different attribute representation levels (*r* = 0.4560, *r* = 0.1410, *r* = 0.2017, respectively, Fig 3a). As V1 had relatively low effective dimensionality (*ED* = 6.829) but high explicit attribute representation level, we conclude that it represents a low-dimensional space including mostly the two explicit attributes. On the other hand, ACC and PCC also represent only a few dimensions (*ED* = 7.759, *ED* = 7.899), but they are probably mostly in a latent space, as their explicit attribute representation levels were low while capturing task-relevant information. This also applies to area MT (*ED* = 5.831, *r* = 0.1868). Finally, EC and vSTR had low attribute representation levels (*r* = 0.1691, *r* = 0.2211), meaning that their high dimensionality (*ED* = 16.04, *ED* = 17.56) consists primarily of latent dimensions. These results build upon the claim that the pre-defined ROIs represent a mixture of explicit and latent attributes and show that they also include a mixture of both high- and low-dimensional latent spaces.

Next, to estimate the relationship between the explicit attribute representation levels and effective dimensionality broadly, we conducted a whole-brain analysis similarly to the pre-defined ROIs. First, we estimated the effective dimensionality for each of 419 parcels and then correlated it with their explicit attribute representation levels, calculated in the previous section. We found a strong significant negative relationship using a robust correlation between the two metrics in both samples (*r*_*first*_ = −0.3045, *r*_*replication*_ = −0.3679, *p* < 10^−8^, Fig 4e), indicating that areas with higher dimensionality showed lower explicit attribute representation levels. This pattern is expected, as greater dimensionality reduces the relative contribution of each dimension.

Finally, we tested whether effective dimensionality improves predictions of the decoy effects. We used the 10,000 combinations of eight randomly selected brain parcels and calculated the average effective dimensionality for each combination. Then, we calculated the robust correlation between the average effective dimensionality and the prediction error of the corresponding RDM regressions. We found a very weak negative correlation between effective dimensionality and prediction error in both the first (*r* = −0.0240, *p* = 0.0162) and the replication samples (*r* = −0.0564, *p* < 10^−7^), indicating that brain parcels with higher dimensionality predict the decoy effect slightly better. As the effect sizes were small, we conclude that dimensionality alone is not a strong predictor of the decoy effect. Together with the pre-defined ROIs results, we conclude that a mixture of high- and low-dimensional representations, representing both explicit and latent attributes, support the prediction of context effects from neural representations.

## 3. Discussion

In this work, we showed the advantages of using high-dimensional neural representations to predict choice context effects. Specifically, we predicted the magnitude of decoy effects elicited by lottery sets as estimated in a large behavioral sample without relying on the lotteries’ explicit attribute space of amount and probability. Instead, we used only the similarity between the lotteries in the neural representation space as estimated in two independent fMRI samples. Furthermore, we showed that these neural representations do not only represent the explicit attribute space, but that higher-level value areas might represent latent dimensions that extend beyond the explicit attribute space, while primary visual areas represent the stimuli’s explicit attribute space to a higher extent, even after controlling for perceptual similarity. Value-related areas also exhibited higher effective dimensionality in their representations, compared to visual areas, which was related to lower levels of explicit attribute representation. Together, our results emphasize the strength of relying on neural representations when analyzing choice behavior, leveraging the entire set of attributes that participants consider without explicitly modelling them.

While we introduced neural representational similarity to the analysis of choice context effects, the concept of similarity is hardly novel in this field. Since Tversky’s similarity hypothesis^5^, many studies have used similarity in explicit attribute space to explain biases in human choices^3,4,8,11,13,14^. Here, we extend this framework to high-dimensional neural spaces, also capturing similarity in latent attributes encoded by the neural system that were previously inaccessible to researchers. This generalized framework offers a neural explanation for the decoy effect, based on the relative positions of options in neural space, regardless of specific systems or stimuli. As such, it could also explain the decoy effects observed across a large variety of animals^33–35^, where the choice options had much more complex attribute spaces than those in typical human studies.

Over the years, several studies have failed to elicit decoy effects using complex choice options such as images of food products^21,23^. However, these attempts faced several methodological criticisms^36,37^. For a decoy options to influence choice, it should be perceived as both similar and inferior to the target. With complex stimuli, this similarity is not necessarily recognized, since participants might infer the attributes of images differently than intended by the researchers. Yet, even a more recent attempt which assessed the similarity between image stimuli and constructed participant-specific target-decoy pairs based on subjective evaluations, has failed to generalize the decoy effect^38^. Nonetheless, the similarity and subjective measurements were analyzed separately and were not combined. Neural representations overcome this limitation by integrating perceptual similarity, value-related information, and their interactions in a single framework. By capturing these properties together, and without requiring pre-defined attributes, this approach holds great promise for generalizing the decoy effect to real-world scenarios using more naturalistic stimuli.

Throughout this work, we used the term “latent attributes” to describe the information encoded in the neural representations but not explained by the lotteries’ two explicit attributes. Although the lotteries were described only by amount and probability, many features can be extracted from them, such as complex utility calculations, perceptual properties, or numerical heuristics that make some numbers easier to process^19,20^. Nonetheless, defining these attributes a priori and using them in a choice model could be extremely hard. Thus, we do not aspire to describe these attributes explicitly, and do not rule out the possibility that they could be captured with sufficiently detailed computational economic and visual models. Instead, our main contribution is to show that the neural representation approach leverages these “latent attributes” without requiring them to be modelled directly.

Leveraging neural data to improve predictions of human decision-making is one of the main goals of the field of neuroeconomics, connecting between neuroscience, economics, and cognitive psychology^39^. Here, we show the advantage of combining concepts from vision neuroscience^16–18^ and behavioral economics^5,8^ to gain access to latent dimensions that are represented internally by the decision-makers and extend beyond the explicit attributes presented to them. These dimensions were inaccessible in previous studies and therefore, neglected, and incorporating them into our models enhanced performance in predicting choice behavior.

Regardless of the contribution to decision-making research, our findings also relate to previous works in vision neuroscience. Specifically, the attribute representation levels analysis revealed that visual areas represent semantic, and not purely perceptual, attributes (Fig 3a-c). V1, as well as other parcels in the occipital lobe, represented the stimuli’s attribute space to a large extent, even when controlling for the perceptual similarity between the stimuli (Fig 3a-e). This is in line with findings describing the top-down influences on activations in visual areas, sometimes referred to as cognitive maps, which combine visual and top-down information^40,41^.

Moreover, we showed that representations of individual stimuli, presented separately, can predict choice behavior when these stimuli are later combined. This relates to a broader discussion on whether neural representations of object combinations reflect a simple summation of the individual components or a context-dependent integration which deviates from this sum^42^. In our experimental design, participants viewed single lotteries during the scan, preventing us from directly measuring the representation of their combinations. Nevertheless, we ascribe our ability to predict the behavioral choice context effects using only the neural similarities between individual lotteries to the estimations of the relative position in neural space of each lottery option in respect to the other lotteries in the choice sets. Even when analyzed in the two-dimensional explicit attribute space, these relative positions can predict the effects of context on choice (as in the explicit attribute model). Remarkably, incorporating both explicit and latent attributes in the neural space leads to superior performance. This suggests that neural similarity conveys enough information about how options interact when presented together, allowing us to predict choice context effects.

To conclude, we showed that neural representations predict human choice context effects, while relaxing the strict assumptions regarding the underlying attribute space of the options. By leveraging neural data, we could access latent dimensions encoded by participants, improving predictions of choice behavior compared to traditional methods. Our general framework is applicable to any type of stimulus and thus could help decision-making research focus on more naturalistic stimuli.

## 4. Methods

### 4.1 Participants

#### 4.1.1 fMRI participants

We recruited a total of 70 participants (43 females, mean age 25.51 ± 3.30), separated into two samples. The first sample included 31 participants (18 females, mean age 26.20 ± 3.51). Two participants were excluded for failing to respond in more than 30% of trials and one participant was excluded due to artifacts in their scans. We therefore reported the analysis of the 28 remaining participants in the first sample. After analyzing this sample, we preregistered our methods and performed a power analysis (https://aspredicted.org/s633-nkb6.pdf), collected and analyzed a replication sample of 39 additional participants (25 females, mean age 24.97 ± 3.02). Four participants were excluded from the replication sample for responding very inconsistently and one participant was excluded due to artifacts in their scans. We therefore reported the data of the remaining 34 participants in the replication sample. All participants gave their informed consent before participating in the study, which was approved by the local ethics committee at Tel-Aviv University and by the Institutional Review Board committee of Sheba Medical Center.

#### 4.1.2 Behavioral participants

We recruited a behavioral sample of 135 participants (79 females, mean age 23.14 ± 4.12). We aimed for a sample size of 120, similar to previous studies investigating between-participant decoy effects with experiments run in-person^9,23^. Thirteen participants were excluded due to failing more than one (out of nine) “catch” trials. We therefore reported the analysis of the remaining 122 participants. All participants gave their informed consent before participating in the study in return for monetary compensation or course credit. The study was approved by the local ethics committee at Tel-Aviv University.

### 4.2 Behavioral tasks

#### 4.2.1 Lottery evaluation task

We used the standard Becker-DeGroot-Marschak (BDM) task^43^, a common incentive-compatible procedure in economic literature to measure participants’ willingness-to-pay. In the task, participants were first given an initial budget of 80 NIS (≈$21). They then saw one lottery on each trial, offering a probability to win a certain amount of money (and otherwise, win nothing), and were asked to choose how much they were willing to pay in order to participate in the lottery. At the end of the task, one trial would be selected at random, and a random price would be determined from 0 NIS and up to the full budget (80 NIS). If the participants’ choice in this trial was lower than the random price, they would not participate in the lottery and would not pay anything, keeping the full budget. If the participants’ choice in this trial was higher than the random price, then participants would pay the determined random price from the budget, participate in the lottery, and keep the remaining budget. Participants viewed a total of 31 unique lotteries which were repeated in a random order in five (fMRI samples) or three blocks (behavioral sample). Participants in the fMRI samples performed this task inside the MRI scanner and had three additional trials with a blank screen in each block. Participants in the behavioral sample performed this task to allow them to have similar familiarity with the stimuli compared to the fMRI participants.

In each trial, the lottery was first presented for 1s, followed by the addition of a slider below the lottery to make the choice. The slider and lottery were presented together for a maximum of 5s (fMRI samples) or 4s (behavioral sample) and disappeared after participants made their choice. The inter-trial-interval (ITI) was 6s (jittered) for the fMRI samples and 1s (jittered) for the behavioral sample. The stimulus-onset-asynchrony (SOA) was 11s (jittered) for the fMRI samples, regardless of participants’ reaction times.

Participants were excluded from this task if they failed to respond to more than 30% of the trials, or if their consistency was low between runs, to verify participants were attentive to the task. Low consistency was defined as an average correlation between their answers across runs or an intraclass correlation coefficient (ICC) below 0.6. All exclusion criteria were preregistered.

#### 4.2.2 Choice task

Participants were randomly assigned to either the *binary* or *trinary* groups. In each trial, participants were asked to choose which lottery they would like to participate in. Participants in the binary group were presented with two lotteries in each trial. Participants in the trinary group were presented with three lotteries in each trial which included the same two lotteries as the binary group with an additional “decoy” option. Participants were informed that at the end of the experiment one trial would be randomly chosen and realized based on their choices. Each participant chose between 27 unique lottery sets and performed three blocks of the task. The task also included three catch trials in each block which had one lottery clearly superior to the other(s), both in probability and amount (first order stochastic dominance). Participants who failed at more than one catch trial were excluded. Importantly, the lottery sets used in this task were constructed using the same 31 lotteries from the lottery evaluation task. Participants in all samples completed this task after the lottery evaluation task. Participants in the fMRI samples completed this task outside the MRI scanner. Participants had no time pressure to complete the task. The ITI was 1s (jittered). The choices of the participants in the fMRI samples were not analyzed in this work, to allow the neural representation analysis to be independent from the behavioral results.

### 4.3 Procedure

Participants in the fMRI samples were instructed about the lottery evaluation task and then read an instruction sheet. Then, they completed a practice block of the task outside the MRI scanner using a trackball. The practice block included five example lotteries not used in the real task and had the same time limits and ITI as the real task. After the practice was completed, participants entered the scanner and performed a resting-state fMRI scan and were asked to stay awake while staring at a fixation cross. Then, participants performed the same practice block, this time inside the scanner, using an MRI-compatible trackball. Participants then performed five blocks of the lottery evaluation task. After completing the scans, participants were instructed about the choice task, which was completed outside the scanner. Participants completed two practice trials, and then completed three blocks of the task. At the end of the experiment, one of the trials was randomly selected from either the lottery evaluation or choice task. Some fMRI participants did not complete the choice task due to tiredness after the scans. The trial was realized for monetary payment based on the instructions of the selected task. The average prize was 58 NIS + 100 NIS show-up fee (≈$42 total).

Participants in the behavioral sample were instructed about both tasks and read two instruction sheets. Then, they completed one practice block and three blocks of the lottery evaluation task, followed by the practice block and three blocks of the choice task. Finally, participants received payment based on one random trial, similar to participants in the fMRI samples. The average prize was 49 NIS (≈$13), with an additional course credit for students or an additional show-up fee of 25 NIS (≈$6.5) for the other participants.

### 4.4 Decoy effect analysis

To calculate the decoy effect for each lottery set, we calculated the difference in target choice ratio between the trinary and binary groups. In detail, we first calculate the relative choice share (RCS) of the target (i.e., number of choices of target divided by the total number of choices of the target and competitor) for each lottery set and each group. Following previous studies, the decoy effect was defined as the difference between the trinary and binary RCS^11,14,44–48^, resulting in 27 decoy effects, one for each unique lottery set in the choice task. The decoy effect was calculated solely based on the behavioral sample, to make it independent from the neural data of the fMRI samples.

### 4.5 Image acquisition

Participants completed a functional Magnetic Resonance Imaging (fMRI) session including anatomical, task fMRI, and resting-state fMRI scans. Scanning was performed at the Strauss Neuroimaging Center at Tel Aviv University, using a 3T Siemens Prisma scanner with a 64-channel Siemens head coil. T1-weighted anatomical images were acquired using a MPRAGE scan (TR = 2400ms, TE = 2.78ms, 0.9mm^3^ voxels). T2-weighted anatomical images were acquired using a SPACE (SPC) sequence (TR = 3200ms, TE = 554ms, 0.9mm^3^ voxels). The fMRI scans were acquired using a T2*-weighted functional multi-band EPI pulse sequence (TR = 750ms, TE = 30.8ms, 2mm^3^ voxels, multi-band factor = 8, 72 slices with no inter-slice gap). All anatomical and functional scans were aligned -30° to the AC–PC plane to reduce signal dropout in the orbitofrontal area. The task fMRI consisted of five runs, each lasting ≈7 minutes. The resting-state scan lasted 10 minutes, and participants were instructed to keep their eyes open.

### 4.6 Preprocessing

We used the Human Connectome Project (HCP) minimal preprocessing pipeline^49^, a series of image processing tools designed by the HCP to remove artifacts and distortions and perform denoising and registration to standard space. To implement the HCP pipeline, we used the Quantitative Neuroimaging Environment & Toolbox (QuNex)^50^. Briefly, the fMRI data underwent the following steps: gradient and EPI distortion corrections, motion correction and nonlinear alignment, and registration to MNI standard space^51–53^. The data was then denoised using FMRIB’s ICA-based Xnoiseifier (FIX)^54,55^, which identified noise and motor components in the functional data and removed them. Finally, the denoised data was resampled onto a surface of 91,282 “grayordinates” in standard space.

### 4.7 Representational Similarity Analysis regression

Our main aim was to predict the decoy effect using high-dimensional neural representations, without relying on the stimuli’s explicit attributes. We leveraged concepts and ideas from the two-dimensional analysis of the decoy effect in the decision-making literature and applied them to neural representations. Specifically, we deployed a regression-based Representational Similarity Analysis (RSA)^17^. Briefly, we extracted neural representations for each lottery, calculated similarity patterns between each pair of lotteries, and used the average similarity patterns across participants to predict the decoy effect that each lottery triad elicited in the behavioral sample. We performed both cross-validated out-of-sample predictions and in-sample data-fitting procedures.

#### 4.7.1 Pre-defined regions of interest

We aimed to use the latent neural representation space of the choice options, which is not limited to their explicit attributes. Therefore, we based our neural representation analysis on several pre-defined regions of interest (ROIs), to include both low-level sensorimotor as well as high-level value-related information.

The value-related ROIs included the anterior cingulate cortex (ACC)^56^, posterior cingulate cortex (PCC), ventromedial prefrontal cortex (vmPFC), and ventral striatum (vSTR), based on previous works showing their activations correlate with choice behavior and utility information^57–59^. Additionally, the entorhinal cortex (EC) was selected based on a recent work showing its activations track the position of options in an abstract value space^60^. The sensorimotor ROIs included the bilateral areas of V1 and MT from the multi-modal atlas^61^ and bilateral M1 based on motor contrasts from the HCP data^62^. V1 and MT were chosen to include the task’s visual information across the visual system hierarchy, as well as value-related signals^63–65^. M1 was chosen due to the substantial motor response required in the task, and following works showing the value-related signals in motor areas^66–70^. Overall, we had eight pre-defined ROIs.

#### 4.7.2 RSA

To compute the neural representational similarity, we used the ubiquitous RSA framework^17^. First, we fitted a General Linear Model (GLM) with 32 regressors, one for each of the 31 lotteries and an additional regressor for the three blank trials. Lottery regressors were defined for the duration of viewing the lottery and before making the choice (1s). For each participant we fitted a GLM for each run, resulting in five GLMs per participant. Next, we calculated the contrast maps for each lottery in each run (zstat maps) and averaged the contrast maps for each participant across runs. Trials with large motion (RelativeRMS>0.75) were excluded from this average. Then, for each participant we extracted the average multivertex pattern for each lottery and for each ROI, and calculated the (squared) Euclidean distance between each pair of patterns for each ROI. Therefore, in the main analysis we had eight 31x31 neural representational dissimilarity matrices (RDMs) for each participant, one per pre-defined ROI.

#### 4.7.3 Neural RDM regressions

Ultimately, the main question in this work referred to the effect of stimuli on the *average* behavior of a group. Accordingly, we planned to use the average neural similarity (i.e., the average brain) from the fMRI samples to predict the group-level choice behavior of the behavioral sample. Therefore, the last step in our analysis pipeline was to use the average neural RDM of each lottery set to predict the decoy effect that this lottery set elicited in the behavioral sample. For each participant, we first normalized each RDM to zero mean and unit variance, and then averaged the ROIs’ RDMs across participants, resulting in eight averaged RDMs.

Finally, we trained regression models that used the average RDMs to predict or fit the decoy effects elicited by each lottery set. Note that we have a decoy effect value for each one of 27 lottery sets, and each lottery set was constructed by 3 lotteries: the target, competitor, and decoy options. Therefore, the RDM relevant for each lottery set is a symmetric 3x3 matrix, describing the neural dissimilarity between the three lotteries of the lottery set. The regression model thus had 3 regressors for each ROI corresponding to the three bottom entries of the 3x3 RDM:

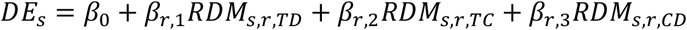

Where *s* is the lottery set *s* ∈ {1, …,27}, *r* is the ROI *r* ∈ {*V*1, *MT, M*1, *vmPFC, ACC, PCC, vSTR, EC*}, *TD* is the target-decoy dissimilarity, *TC* is the target-competitor dissimilarity, and *CD* is the competitor-decoy dissimilarity.

##### 4.7.3.1 Out of sample predictions

To predict the decoy effects out of sample, we trained a Lasso regression model and evaluated it using leave-one-lottery-out cross-validation. As some lottery sets had overlapping lotteries (e.g., the target in set #12 is the competitor in set #27), in each fold we selected one lottery and held out all lottery sets that contained it. Overall, we had 25 unique folds each with between 3-8 lottery sets held out. In each fold, we performed hyperparameter tuning to select the regularization factor, using an inner leave-one-lottery-out within the training set. Finally, we evaluated the models by their average root mean squared error (RMSE) across the folds, and by their average Spearman correlation between predicted and actual values across the folds. To estimate the significance of the prediction performance, we did a permutation test by shuffling the decoy effects and training and evaluating the models on the shuffled data (*n* = 10,000 permutations).

To explore the distribution of predictive power across brain areas beyond our pre-defined ROIs, we performed an exploratory whole-brain analysis. First, we parcellated the brain to 419 parcels using a whole-brain parcellation^28,29^ and calculated the RDMs for each parcel. Then, we selected eight random parcels and trained and evaluated Lasso regression models to predict the decoy effects (*n* = 10,000 combinations). We then calculated how many combinations performed better than our eight pre-defined ROIs.

##### 4.7.3.2 Data fitting

As we had too many regressors compared to the dataset size when using all eight pre-defined ROIs, we used a forward stepwise regression procedure to select a subset of these ROIs. We deployed this stepwise procedure while ensuring that all three regressors of each ROI would be included (or removed) together, to preserve the interpretability of our models. We deployed this procedure only on the RDMs of the first sample and then used the selected ROIs in the replication sample using standard linear regression. We evaluated the models using their 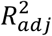 scores and by the correlation between fitted and actual values. To account for the feature selection procedure, we corrected the p-value of the forward stepwise regression for all the models tested using False Discovery Rate^71^. The forward stepwise regression was applied only on the first fMRI sample to select ROIs. For the replication sample, we fitted a standard linear regression using the set of ROIs selected in the first sample.

### 4.8 Attributes-based models

To establish a baseline for comparing the neural RDM regression predictions, we fitted regression models based on the stimuli’s explicit attributes and participants’ WTP responses. Similar to the RDM models, we first used each model to train and evaluate a Lasso regression model to predict the decoy effects in cross-validation, and calculated its significance using a permutation test (*n* = 10,000 permutations). We then also fitted a standard linear regression model to the decoy effects and calculated its 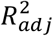 and correlation between fitted and actual values.

#### 4.8.1 Explicit attributes models

First, we used the lotteries’ raw attributes of amount and probability resulting in six regressors:

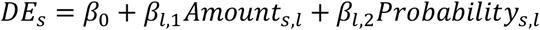

Where *l* ∈ {*Target, Competitor, Decoy*}, and lottery set *s* ∈ {1, …,27}.

We also trained variations to this model including the lotteries’ expected values (EVs):

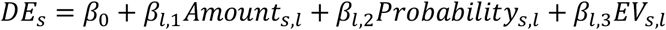

#### 4.8.2 WTP models

Next, we trained WTP-based models, using the participants’ average WTP response for each lottery. As all three samples performed the lottery evaluation task, we trained three models, one using the average WTP response from each sample. Each model included three regressors:

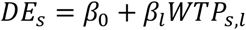

Additionally, we trained a model including the difference between the average WTP response of each pair of lotteries:

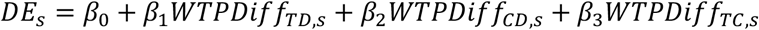

Finally, we combined the explicit attributes with the WTP values in a single model:

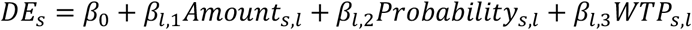

### 4.9 Computational modelling

To verify our behavioral results are aligned with previous studies exploring the decoy effect, we fitted several computational models used to explain the decoy effect. In contrast to the other models used in this work, these models are not regression based. Instead, they are used to fit participants’ individual choices, rather than the aggregated decoy effects directly.

To fit the models, we first scaled the amounts and probabilities of the lotteries to be between zero and one using min-max scaling. Then, we fitted the model’s parameters to all the participants’ choices using the Nelder-Mead simplex algorithm implemented by *scipy*’s *fmin* function^72^ or MATLAB’s *fminsearch*. Next, we calculated the predictions for the probability to choose the target option for each triad when given all three lotteries (including the decoy option) and when given only two lotteries (without the decoy option). Finally, we calculated the difference between these two predictions to get the predicted decoy effect. Similar to the regression models, we trained and evaluated the models using cross-validation. We then also fitted the model to the entire choice data and evaluated the in-sample predictions using the correlation between actual and fitted decoy effects. The computational models were not tested for their significance level due to their long optimization procedures.

#### 4.9.1 Normalization-based models

First, we used the Divisive Normalization model^14,24^, which estimates each option’s subjective value by its raw value divided by the sum of all options’ values:

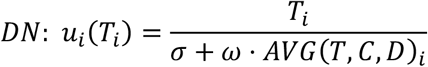

Where *i* is the attribute (amount or probability), *u*_*i*_(·) is the utility function, *T, D, C* are the target, decoy, and competitor. *σ* and *ω* are free parameters representing the bias and weight of the normalization procedure.

Next, we used the Recurrent Divisive Normalization^14,24^, overweighing the value of the estimated option:

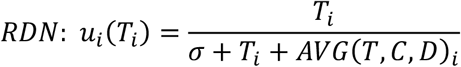

Finally, we used the Adaptive Gain model which assumes the utility of each choice option is dependent on the context via a sigmoidal nonlinearity:

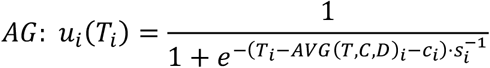

Where *s*_*i*_ is the sigmoid slope and *c*_*i*_ is the sigmoid bias (together with the average attribute of *T, D, C*) of attribute *i*.

To form each option’s final utility, for all models we aggregated the utilities by weighing each attribute by a weight parameter *w*:

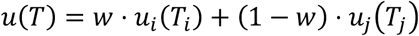

Finally, the choice was made by a softmax function with temperature parameter *τ*:

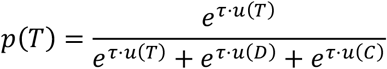

#### 4.9.2 Evidence accumulation (drift-diffusion) models

Second, we used dynamic models based on evidence accumulation (drift-diffusion), which have previously shown to explain the decoy effect. Namely, we used the Mutual Inhibition model^27^, incorporating an inhibition mechanism between the accumulating neurons, and the Selective Integration model^25,26^, incorporating an attentional shift mechanism favoring the currently-better option.

Following previous implementations, both models were trained only on the target and competitor choices, excluding the small portion of decoy choices (3%). The Selective Integration was jointly trained to fit participants’ choices and reaction times, following the procedure described by Cao & Tsetsos^26^. The Mutual Inhibition model was run for 1000 iterations to generate choice probabilities, following the procedure described by Chau et al.^27^.

### 4.10 Explicit attribute representation analysis

To measure the extent to which each ROI represents the lotteries’ explicit attributes of amount and probability, we first calculated the explicit attribute RDM. To do so, we calculated the (squared) Euclidean distance between each pair of lotteries based on their amount and probability values, resulting in a 31x31 RDM. Next, we correlated the bottom triangle of the neural RDM of each ROI with the bottom triangle of the explicit attribute RDM to estimate the ROI’s attribute representation level. We performed this procedure for each of our eight pre-defined ROIs.

Although the position in the lotteries’ two-dimensional explicit attribute space relies only on the amount and probability values, the distance between lotteries also contains perceptual information (e.g., 10 and 11 are close both perceptually and in value). To account for this, we also estimated the attribute representation levels after controlling for the perceptual similarity between the lotteries. To do so, we first extracted the activations from the first layer of a pretrained convolutional neural network^30^ for each lottery stimulus (represented as an image), and calculated the (squared) Euclidean distance between each pair of activations to form a perceptual RDM. We then calculated the partial correlation between the explicit attribute RDM and the neural RDM, regressing out the perceptual RDM.

#### 4.10.1 Whole-brain attribute representation levels

To look at the explicit attribute representation levels at a wider lens, we performed the same procedure for each brain parcel out of 419 in the whole-brain exploratory analysis (see *Out of sample prediction* section). After projecting the resulting attribute representation levels to a brain surface, it seemed there is a posterior-anterior quadratic gradient to the values. To test this, we fitted a quadratic function to take the posterior-anterior coordinate and explain the attribute representation level.

### 4.11 Effective dimensionality analysis

To explore the latent space of neural representations, we estimated the effective number of dimensions represented in each ROI. We used the effective dimensionality metric^31,32^ which is a continuous measure quantifying the unique information represented in a set of measurements, using entropy-based metrics.

To calculate the effective dimensionality of each ROI, we created a representation matrix for each participant containing the multivertex pattern for each lottery. Each representation matrix is in ℝ^*KxV*^, where *K* = 31 is the number of lotteries, and *V* is the number of vertices in the ROI. Next, we calculated the covariance matrix for the representations and performed eigenvalue decomposition. Then, we calculated an entropy-based metric that corresponds to the number of uncorrelated measurements in the data, based on the eigenvalues^31^:

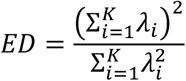

Where *λ*_*i*_ is the *i*-th eigenvalue of the covariance matrix. *ED* can range between 1 to *K* = 31. Lower values of *ED* indicate that the lotteries’ representations are described by a small number of dimensions, and higher values indicate that the representations are described by a large number of dimensions.

## Data Availability

The processed fMRI data, raw behavioral data, and stimuli are available at https://github.com/asafmm/neuro_decoy_effect.

## Code Availability

All analyses codes are available at https://github.com/asafmm/neuro_decoy_effect.

## Acknowledgements

The authors acknowledge with thanks the support of the Israel Science Foundation (1432/23) and the Henry Crown Institute of Business Research in Israel.

## Author contributions

A.M., I.T., and D.J.L. conceived the project. A.M. designed the experiments, collected the data, and performed the analyses. A.M. and T.Z. collected the fMRI data. A.M., I.T., and D.J.L. wrote and edited the paper.

