## Supplementary materials for "Neural similarity between choice options predicts group-level context effects"

### Supplementary Results and Methods

#### 1. Choice probabilities are explained by decoy-target similarity and inferiority

We explored the effects of the choice options attributes on the probability to choose the target option separately for the binary and trinary groups. We used the choice options' attributes (amount and probability), the differences between their expected values (EV), and the difference between their willingness-to-pay (WTP) calculated individually for each participant based on their evaluations in the evaluation task. All regressors were normalized before fitting. All results are reported in Table S1. The model for the trinary group:

$$\text{logit}\, P\left(T_p = 1\right) = \beta_0 + \beta_{1,l} Amount_l + \beta_{2,l} Probability_l + \beta_3 (EV_T - EV_C) + \beta_4 (EV_T - EV_D) + \beta_5 (WTP_T - WTP_C) + \beta_6 (WTP_T - WTP_D) + \beta_7 RT + \sigma_p$$

Where $l \in \{T, C, D\}$, representing the Target, Competitor, and Decoy, respectively, WTP is the willingness-to-pay value based on participant $p$'s responses in the evaluation task. $\sigma_p$ is a participant-level random intercept. For the binary group, we used the same model excluding the regressors related to the decoy option.

First, we found that participants in both binary and trinary groups responded according to the EV differences. That is, the probability to choose the target option increased with higher target-competitor EV difference ($\beta_{trinary} = 1.06$, $\beta_{binary} = 0.9$, $ps < 0.001$). The WTP difference showed the same effect ($\beta_{trinary} = 0.51, \beta_{binary} = 0.17, ps < 0.001$), suggesting that participants generally responded according to their previous evaluations.

Moreover, we found that in the trinary group, the probability to choose the target option also increased when the target and decoy's EVs were closer ($\beta = -0.24, p < 0.001$), in line with previous findings emphasizing the target's similarity and domination over the decoy driving the choice to the target[1]. Similarly, lower monetary amount of the decoy option predicted higher probability to choose the target option ($\beta_{DecoyAmount} = -0.47, p < 0.001$), while the decoy's probability attribute had no significant effect on choosing the target option ($\beta_{DecoyProbability} = -0.07, p = 0.22$), showing that participants were affected more by the amount than by the probability attribute.

**Table S1.** Estimates of a mixed-effects linear model of the probability to choose the target option in each group.

| | Trinary group | Binary group |
|---|---|---|
| | Estimate (Std. Error) | Estimate (Std. Error) |
| Intercept | -0.09 (0.05) | -0.21*** (0.05) |
| Target amount | -0.59*** (0.11) | -0.45*** (0.08) |
| Target probability | 0.64*** (0.13) | 0.51*** (0.10) |
| Competitor amount | 0.50*** (0.17) | -0.11 (0.16) |
| Competitor probability | 0.13*** (0.07) | 0.008 (0.06) |
| Decoy amount | -0.47*** (0.09) | |
| Decoy probability | -0.07 (0.06) | |
| Target-Competitor EV difference | 1.06*** (0.10) | 0.90*** (0.10) |

| | | |
|---|---|---|
| Target-Decoy EV difference | -0.24***<br>(0.09) | |
| Target-Competitor WTP difference | 0.51***<br>(0.05) | 0.17***<br>(0.04) |
| Target-Decoy WTP difference | -0.04<br>(0.05) | |
| RT | 0.02<br>(0.03) | -0.01<br>(0.04) |
| Pseudo-$R^2$ | 0.27 | 0.23 |
| Individuals | 62 | 60 |
| Observations | 5016 | 4797 |

*Note*. The model also includes random effects at the subject level. All regressors were normalized before fitting. * $p < 0.05$, ** $p < 0.01$, *** $p < 0.001$.

**2. Results for attributes regression models**

To establish a baseline for comparing the neural RDM regression predictions, we fitted regression models based on the stimuli's explicit attributes and participants' WTP responses (Table S2).

First, we calculated the average WTP for each lottery across participants in the behavioral sample, and then trained models to predict the decoy effects based on these WTP values, and found relatively low performance ($RMSE_{WTP} = 0.0889$). We also tried using the average WTP from the first and replication fMRI samples, instead of the behavioral sample, and found similar results ($RMSE_{first} = 0.0852, RMSE_{replication} = 0.0902$).

Next, instead of using the WTP values, we calculated the difference in WTP between each pair of lotteries. We then used the WTP differences to predict the decoy effects but again found relatively low performance ($RMSE_{WTP\_Diff} = 0.0811$).

Then, we used the lotteries' explicit attributes, reaching higher prediction accuracy ($RMSE_{explicit} = 0.0764$). We also tried to combine the explicit and WTP models, by training a new model on both the explicit attributes and participants' WTP, performing better than the WTP models but still worse than the explicit attributes model ($RMSE_{explicit+WTP} = 0.0778$).

Finally, we tried adding the lotteries' EVs to the explicit attributes model, instead of the WTP values. The combined model performed better than the explicit attributes model ($RMSE_{explicit+EV} = 0.0753$) and resulted in the best baseline model.

**Table S2**. Different baseline models using the lotteries' explicit attributes, willingness-to-pay (WTP), and expected values (EVs), to predict the decoy effects.

| Model name | Out-of-sample $RMSE$ |
|---|---|
| Explicit attributes | 0.0764 |
| WTP | 0.0889 |
| WTP differences | 0.0811 |
| Explicit attributes + WTP | 0.0778 |
| **Explicit attributes + EV** | **0.0753** |

**3. Previously suggested computational models of the decoy effect**

We trained several computational models used in previous studies to provide a more robust baseline for comparison. Specifically, we trained the Mutual Inhibition[2,3] and Selective Integration[2,4] drift-diffusion models, which fit choices via an evidence accumulation process with an additional inhibition or attentional shift mechanisms, dependent on the choice options' values. Additionally, we trained Divisive Normalization models[5,6] which, as their name suggests, fit choices via different attribute normalizations. These models have been previously shown to fit choice data and explain the decoy effect, mostly in within-subject designs (as opposed to our between-subject design).

Among all computational models, the best for out-of-sample prediction was the Recursive Divisive Normalization ($RMSE = 0.0824$), while for in-sample fitting the Selective Integration model performed very well ($r = 0.74$). Nonetheless, our RDM models still performed best both in out-of-sample and in-sample predictions ($RMSE = 0.0656,\ r = 0.93$). See Table S3 for the full description.

Based on these results, we concluded that the behavioral patterns in our data are fitted relatively well by the inhibition and attention mechanisms modelled by the drift-diffusion models, in line with previous works.

**Table S3**. Models' performance for out-of-sample and in-sample predictions of the decoy effect. Computational models on the top rows, regression models on the bottom rows. The best performing models in each category are underlined.

| Model name | Out-of-sample $RMSE$ | In-sample $r$ |
|---|---|---|
| Mutual Inhibition | 0.0881 | 0.58 |
| Selective Integration | 0.0969 | <u>0.74</u> |
| Divisive Normalization | 0.1454 | -0.14 |
| Recurrent Divisive Normalization | <u>0.0824</u> | 0.06 |
| Adaptive Gain | 0.0936 | 0.30 |
| Explicit attributes | 0.0753 | 0.81 |
| First RDM | <u>0.0656</u> | <u>0.93</u> |
| Replication RDM | 0.0659 | 0.87 |

**4. Univariate responses encode for subjective value but do not predict the decoy effect better than the RDM models**

It is known that univariate fMRI responses in value-related areas correlate with utility information[7–9]. Here, we aimed to replicate this finding in our dataset in a way that would also enable us to predict the decoy effects between the samples. Thus, for each participant we first calculated the average response for each lottery in each ROI. Then, we normalized the average lottery responses within each participant and averaged the normalized response across participants, to reach an average univariate response for each lottery in each ROI.

Then, to verify these univariate responses are indeed task relevant, we correlated them with the fMRI participants' average WTP responses (Table S4). We repeated this analysis separately for each fMRI sample. As expected, we found most pre-defined ROIs showed highly positive correlations with the WTP values, a result replicated in both fMRI samples, and consistent with previous findings showing these regions encode for subjective value.

**Table S4.** Correlations between univariate ROI responses and average WTP.

| Region | First sample $r$ | Replication sample $r$ |
|---|---|---|
| vmPFC | 0.36* | 0.56** |
| vSTR | 0.77*** | 0.73*** |

| | | |
|---|---|---|
| PCC | 0.10 | 0.11 |
| ACC | 0.50** | 0.37* |
| M1 | 0.37* | 0.56** |
| V1 | 0.56** | 0.75*** |
| EC | -0.13 | 0.13 |
| MT | 0.24 | 0.05 |

*Note.* * $p < 0.05$, ** $p < 0.01$, *** $p < 0.001$

Next, we could use these univariate responses to predict the decoy effects observed in the behavioral sample, in a similar procedure to the other regression models described in the main text. We trained a regression model using the univariate responses of the first fMRI sample, and found it did not predict the decoy effects better than the explicit attributes or RDM models ($RMSE_{univariate} = 0.0830$, $RMSE_{explicit} = 0.0753, RMSE_{RDM} = 0.0656$). For the replication sample, the univariate model performed better than the explicit attribute model ($RMSE_{univariate} = 0.0729$), but not the RDM model ($RMSE_{RDM} = 0.0659$). Notably, in both samples, the univariate models outperformed the WTP-based model ($RMSE_{WTP} = 0.0889$), suggesting the univariate responses contain more relevant information for predicting the decoy effect than WTP alone.

Thus, we showed the univariate brain responses in our pre-defined ROIs highly correlate with participants' WTP, encoding utility information. However, these signals do not perform very well in out-of-sample prediction of the decoy effects, suggesting that our results based on neural representations relied on other representations in addition to utility encoding.

### Supplementary Figures

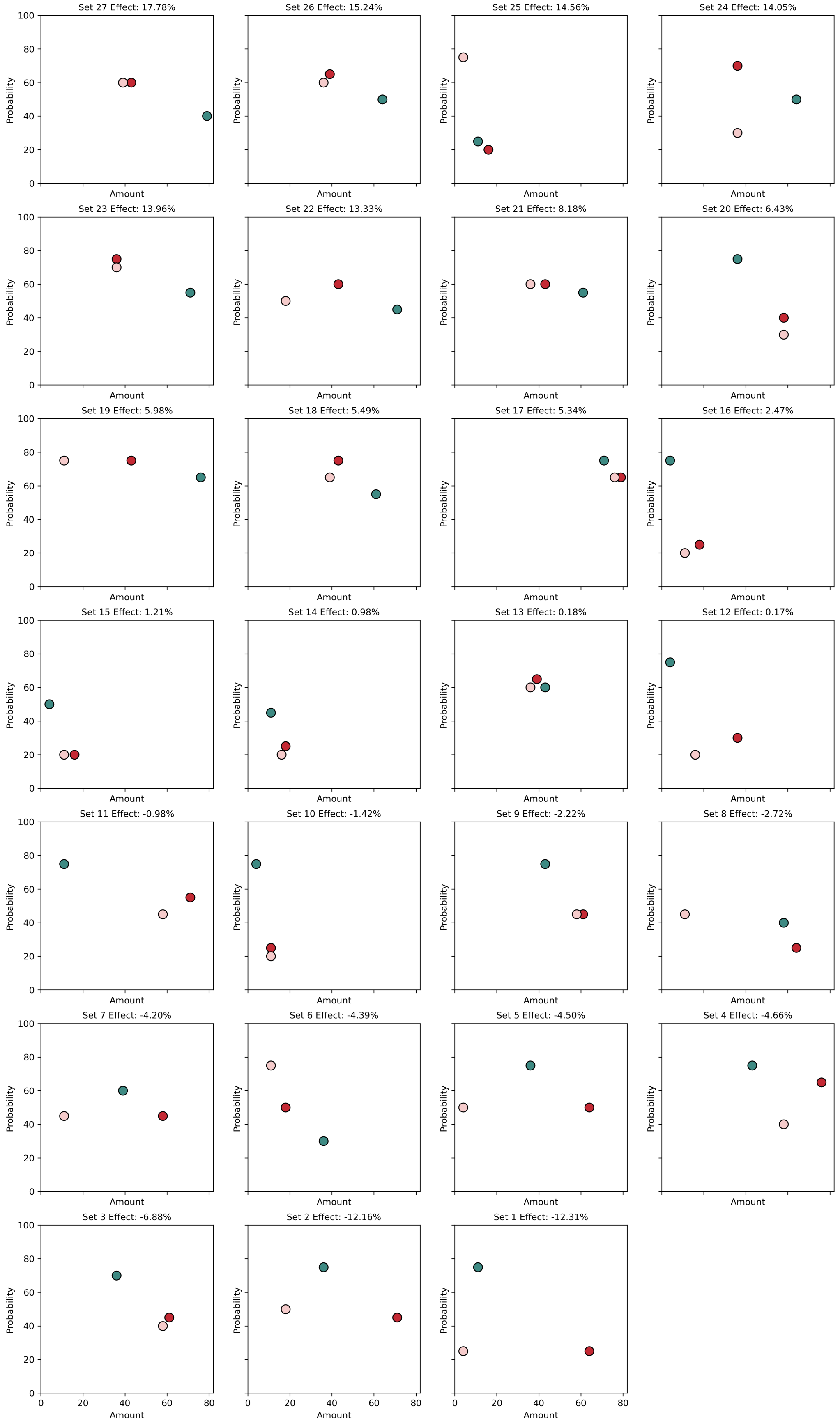


**Figure S1. Lottery sets geometry in the explicit attribute space.** All 27 lottery sets used in the choice task plotted in the two-dimensional explicit attribute space of amount and probability. Color coding similar to Figure 1.

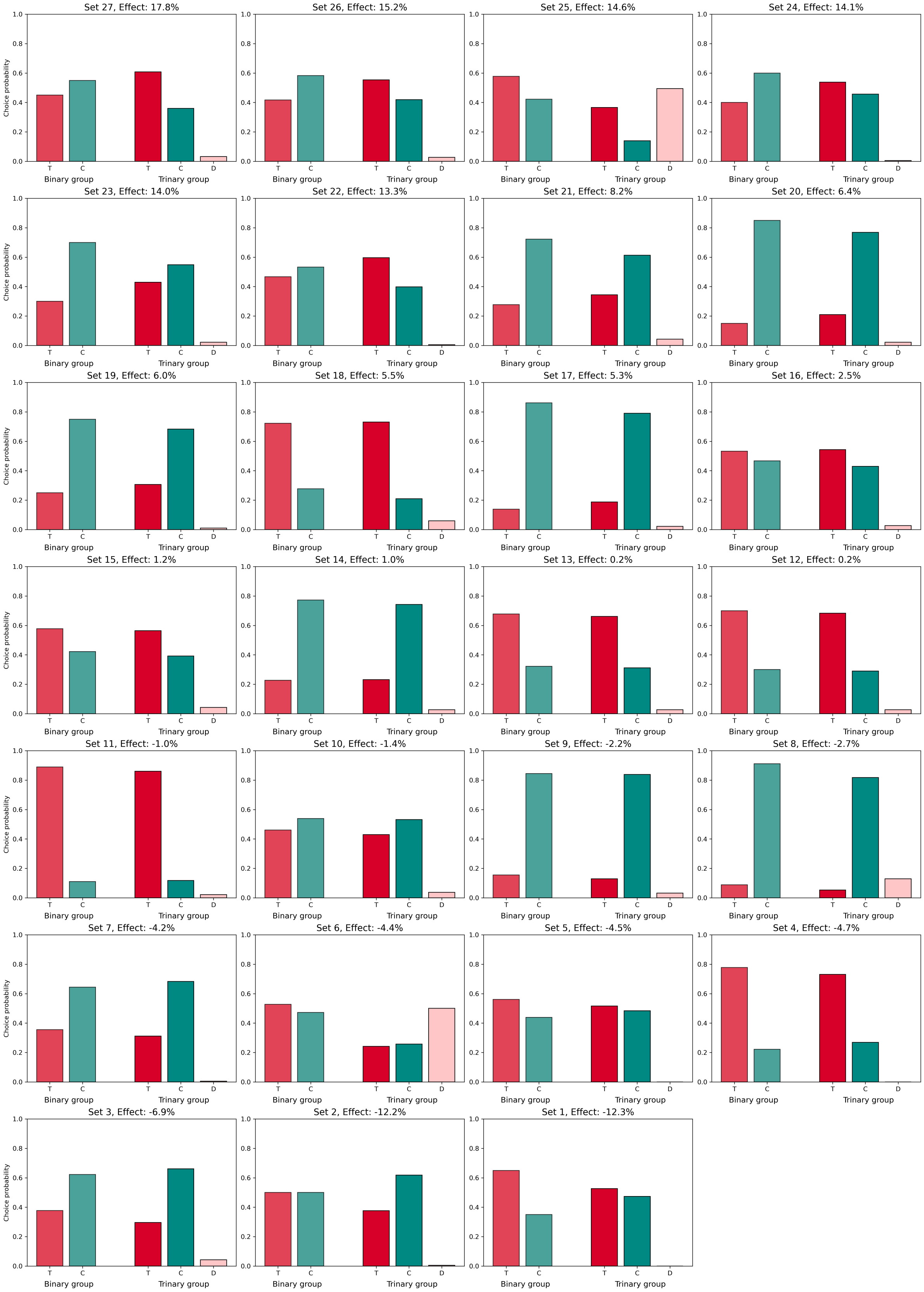


**Figure S2. Absolute choice probabilities for each lottery set.** The target, competitor, and decoy choice probabilities for the binary (left) and trinary (right) groups for each lottery set.

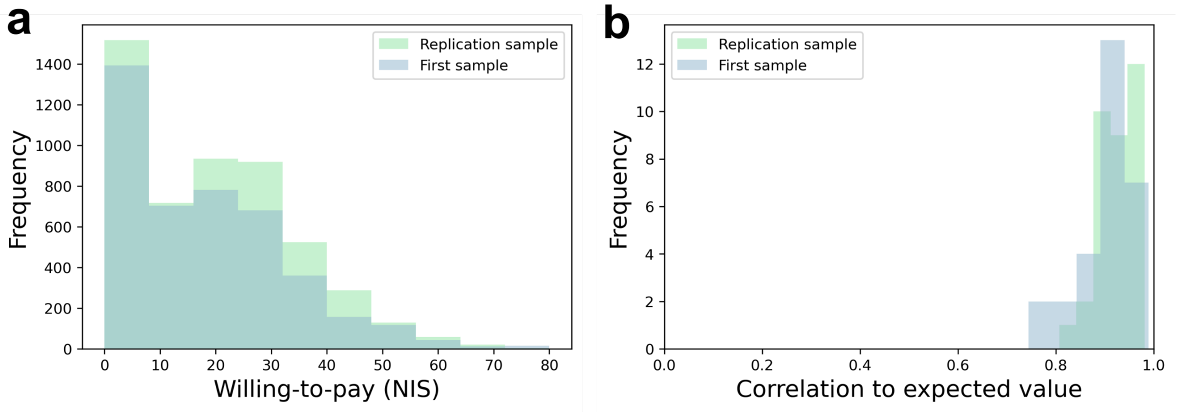


**Figure S3. Evaluation task results from the fMRI samples.** Similar distributions of behavioral results between participants in the first and replication fMRI samples. (a) Distribution of willingness-to-pay of all participants' choices. (b) Distribution of the correlation between willingness-to-pay to participate in each lottery and the lottery's expected value. High correlations show participants understood the task and were attentive.

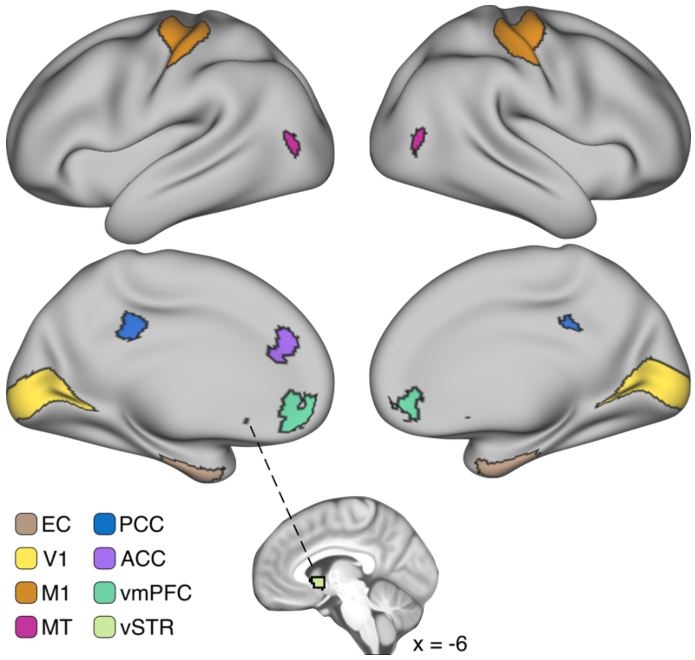


**Figure S4.** The pre-defined ROIs. EC – Entorhinal Cortex, V1 – primary visual cortex, M1 – primary motor cortex, MT – middle temporal visual area, PCC – posterior cingulate cortex, ACC – anterior cingulate cortex, vmPFC – ventromedial prefrontal cortex, vSTR – ventral striatum.

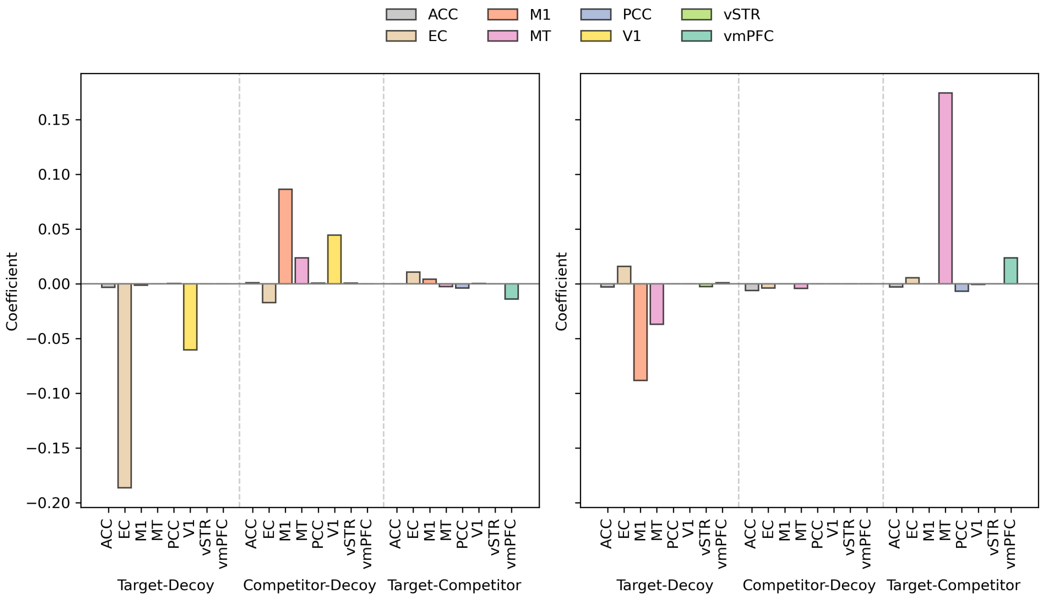


**Figure S5**. Lasso regression coefficients for the first (left) and replication (right) models. Note that the coefficients are in respect to the dissimilarity (not similarity) values.

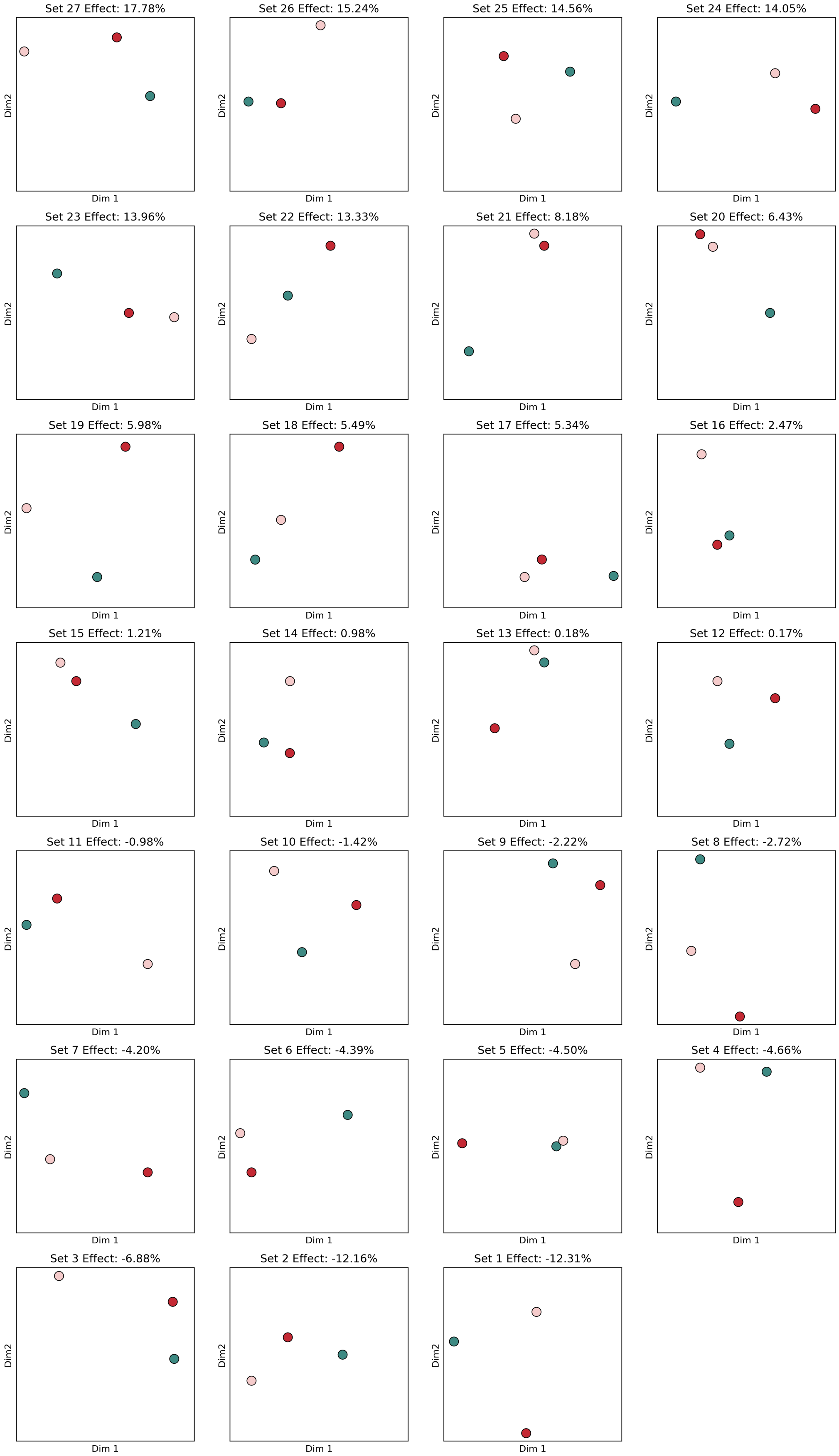


**Figure S6. Lottery sets geometry in neural space.** Non-metric multidimensional scaling visualizations of the EC RDM from the first sample. Visualization of the EC RDM from the replication sample, and from other ROIs, showed similar patterns.

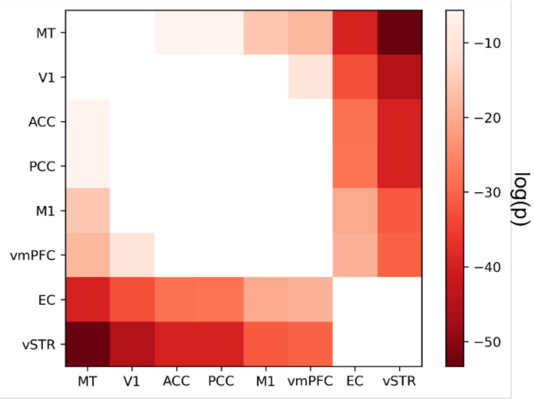


**Figure S7. Pairwise comparisons between the ROIs' effective dimensionality.** The log of the FDR corrected p-values are shown, thresholded at $p < 0.05$ in both the first and replication samples.
